# Combining high resolution and exact calibration to boost statistical power: A well-calibrated score function for high-resolution MS2 data

**DOI:** 10.1101/290858

**Authors:** Andy Lin, J. Jeffry Howbert, William Stafford Noble

## Abstract

To achieve accurate assignment of peptide sequences to observed fragmentation spectra, a shotgun proteomics database search tool must make good use of the very high resolution information produced by state-of-the-art mass spectrometers. However, making use of this information while also ensuring that the search engine’s scores are well calibrated—i.e., that the score assigned to one spectrum can be meaningfully compared to the score assigned to a different spectrum—has proven to be challenging. Here, we describe a database search score function, the “residue evidence” (res-ev) score, that achieves both of these goals simultaneously. We also demonstrate how to combine calibrated res-ev scores with calibrated XCorr scores to produce a “combined p-value” score function. We provide a benchmark consisting of four mass spectrometry data sets, which we use to compare the combined p-value to the score functions used by several existing search engines. Our results suggest that the combined p-value achieves state-of-the-art performance, generally outperforming MS Amanda and Morpheus and performing comparably to MS-GF+. The res-ev and combined p-value score functions are freely available as part of the Tide search engine in the Crux mass spectrometry toolkit (http://crux.ms).

## 1 Introduction

In the analysis of protein tandem mass spectrometry data produced in a bottom-up fashion using traditional, data-dependent acquisition, the database search step is critical. In this step, each observed spectrum is assigned to a peptide sequence drawn from a given database, and the resulting peptide-spectrum match (PSM) is assigned a score. Ideally, a “good” score implies that the peptide was likely to have been responsible for generating the observed spectrum. Of course, in the context of a typical scientific study, the end goal is usually downstream of the PSMs—e.g., to detect and quantify proteins, or to characterize proteins whose quantification changes across experimental conditions. However, none of these downstream steps can be accomplished if the database search step fails. Furthermore, although in principle *de novo* approaches can help to identify some observed spectra, in practice *de novo* approaches do not approach the power of database search strategies to detect hundreds or thousands of peptides in a given complex mixture (1). Consequently, a database search engine forms the backbone of most shotgun proteomics analysis pipelines.

Given the importance of database search, it is not surprising that dozens of search engines have been developed since the advent of the first such search tool, SEQUEST, in 1994 (2) (reviewed by (3)). The algorithms employed by all of these engines are remarkably consistent. For each spectrum, the algorithm extracts from the given database all peptides whose masses fall within a user-specified tolerance of the inferred precursor mass associated with the observed spectrum. Each of these candidate peptides is then scored against the observed spectrum, and the top-scoring peptide is reported as a PSM. Thus, in practice, the defining characteristic of any search engines lies in the details of its peptide-to-spectrum score function. The history of development of shotgun proteomics search engines can be seen primarily as a history of development of PSM score functions.

Some new score functions are driven by technology. For example, over the past decade, the resolution at which tandem mass spectra can be efficiently collected has improved dramatically. Typical data sets offer fragment ion resolution in the range of 5–10 ppm, compared to the ~1 Da resolution that was common a decade ago. This improved resolution means that score functions designed for low resolution data did not necessarily generalize well to higher resolutions. For example, the score function in Morpheus (4) was explicitly designed to make good use of high-resolution mass accuracy.

On the other hand, some new score functions are driven by conceptual advances. A notable trend was the introduction of score functions that aim to achieve good calibration (5–7). We say that a score function is *calibrated* if a score of *x* assigned to one spectrum has the same meaning or significance as a score of *x* assigned to a different spectrum. In practice, many PSM score functions are not well calibrated with respect to spectra, that is, they tend to assign systematically different scores to different spectra. Therefore, performing database search with such a function yields a loss of statistical power (8). One way to improve the calibration of a given score function is to compute, for a given spectrum, the distribution of scores for all possible peptides. The cumulative density function of the resulting distribution then provides a well calibrated score, called a p-value. MS-GF+ demonstrated how to carry out this style of calibration using a dynamic programming procedure (5), and a similar approach was adopted subsequently by RAId aPS (6) and Tide (7).

Unfortunately, a signficant drawback to the dynamic programming calibration procedure is that it typically breaks down when employed in conjunction with data generated using high-resolution fragment mass accuracy. To explain the problem, it is first necessary to outline how the dynamic programming algorithm works. Any dynamic programming procedure involves building up a table of solutions to problems of increasing size. In the case of the procedures employed by MS-GF+, RAId aPS, and Tide, the entry in row *i* and column *j* of the dynamic programming table contains a count of the total number of peptide sequences whose (discretized) mass is *i* and whose (discretized) score with respect to the current spectrum is *j*. The procedure works by filling in values in this table for increasingly large values of *i* and *j*, computing each new value by summing over existing entries in the table (see Methods for details).

The problem arises in the discretization of the mass axis. The logic associated with filling in the table requires sequentially adding together discretized amino acid masses. For this sequential summation to work properly, it must be the case that, for any given peptide, the sum of the discretized amino acid masses equals the discretized sum of the amino acid masses. More concretely, using a bin size of *w* and for a peptide consisting of amino acids *a*_1_*, a*_2_, …, *a_n_*, it must be the case that

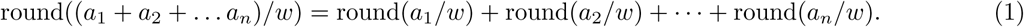

Whether and how frequently Equation 1 is violated depends upon properties of the amino acid masses and the size of the bins used to discretize the mass axis. It is easy to see that, for arbitrary amino acid masses and peptides of reasonable length, Equation 1 will frequently be violated (“Random” series in Figure 1). On the other hand, when we use real amino acid masses and a bin size of ~1 Da, short peptides uniformly obey Equation 1. This is because peptide masses are naturally discrete. Longer peptides do occasionally break the rules because peptide masses are not perfectly discrete; however, for peptides of the size typically considered by shotgun proteomics, this rule-breaking is quite rare. The story changes, however, when we modify the bin size used to discretize the fragment m/z axis. Because such bins do not align well with the natural discreteness of the peptide mass axis, Equation 1 is violated quite frequently. Consequently, the dynamic programming procedure ends up spreading the counts associated with peptides of mass *i* among bins in rows *i*− 1, *i*, and *i* + 1. We are thus left with a conundrum: we have two techniques to achieve improved statistical power—using increased resolution on the fragment m/z axis or using dynamic programming to achieve good score calibration—but we cannot use both of these techniques simultaneously.

**Figure 1:**
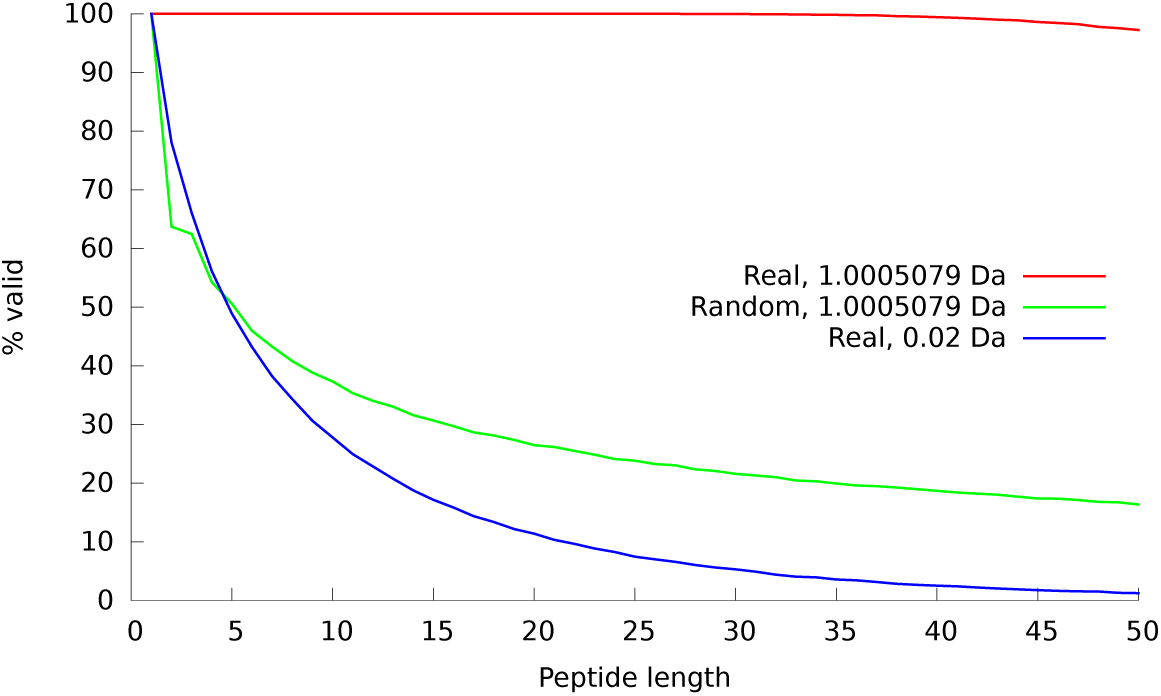
The figure plots, as a function of peptide length, the proportion of randomly generated peptide sequences that obey Equation 1. The two series marked “Real” use monoisotopic masses from the 20 real amino acids; the series marked “Random” uses masses that have a random number in the range 〈0, 1] added to each mass. Two bin sizes (1.005079 Da and 0.02 Da) are used. For each series, a total of 100,000 peptides were simulated.

One solution to this problem is to modify the score function. MS-GF+ does this by creating a score function that takes into account both the intensity of each observed peak as well as its participation in a pair of peaks with a mass difference equal to the mass of an amino acid. The latter term, which is implemented as a weight associated with a given peak, can be computed in high resolution, even if the peaks themselves are scored in low resolution. The dynamic programming procedure can then be carried out in the usual fashion, simply by incorporating these weights.

In this work, we propose an alternative solution. We begin with the XCorr score function, which was included in the very first search engine, SEQUEST, and continues to be used in SEQUEST and a variety of other search engines, including Comet (9), Tide (10), and RAId aPS (6). However, rather than modifying the XCorr score function to take into account high-resolution mass information, we create a new score function, the “residue-evidence” (res-ev) score, that considers pairs of peaks, similar to the MS-GF+ approach. We then score each observed spectrum twice: once with the low-resolution XCorr score that focuses on individual peaks, and once with the high-resolution score that focuses on pairs of peaks. We use dynamic programming to convert each of these scores to p-values, and we employ a previously described method to estimate the p-value for the product of dependent p-values (11).

In this work, we demonstrate that the new res-ev score function provides improved performance on some high-resolution data sets, and that the res-ev and XCorr score functions are complementary to one another. Finally, we demonstrate that the combined XCorr+res-ev p-value yields state-of-the-art performance across a variety of data sets, outperforming MS Amanda and Morpheus and performing comparably to MS-GF+, despite having no trainable parameters. The combined p-value and res-ev p-value score functions are available in the Tide search engine, which is part of the Crux toolkit (http://crux.ms).

## 2 Methods

### 2.1 The XCorr score

The XCorr score function was first described in 1994 as part of the SEQUEST search engine (2) and is still in use today in the commercial SEQUEST product from Thermo Scientific as well as search engines such as Comet (9), Tide (10), X!Tandem (12) and RAId aPS (6).

In our implementation, the first part of the XCorr score involves preprocessing the observed spectrum in four steps, as follows:

1. The mass axis is discretized by creating a vector *O* of mass bins with bin width = 1.0005079 Da. Each bin *O*_*i*_ is assigned an intensity value, which is the maximum of the intensities of observed peaks whose masses fall within the mass range of *O*_*i*_. The bin width is chosen to match the natural quasi-integer masses of peptides and peptide fragments, which in turn derive from the quasi-integer masses of the primary constituent elements of peptides (C, H, N, O, S).
2. The intensity of each bin *O*_*i*_ is replaced by its square root.
3. Peaks with intensity less than 5.0% of the maximum intensity peak are eliminated from the spectrum.
4. *O* is divided into 10 equal length segments, and the intensities within each segment are normalized so the maximum intensity in the segment is 50.
5. A scaled version of *O* is subtracted from itself at each position across a defined window of offsets: 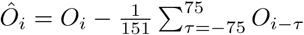

Next, the preprocessed observed spectrum 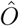 is converted to an “evidence vector” *E* with *n* discretized mass bins, where *n* is the integer mass of the spectrum precursor. Each bin *E*_*i*_ specifies the cumulative evidence for cleavage at some hypothetical position on the backbone of the precursor peptide. More precisely, *E*_*i*_ holds the weighted sum of all intensities in 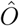 whose mass is consistent with a cleavage producing a b ion with integer mass *m*_*b*_ = *i*:

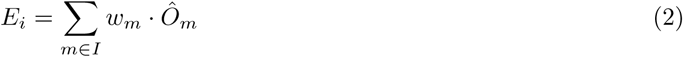

where *I* are the integer masses of the b, y, and neutral loss ions consistent with *m*_*b*_ and *n*, and *w*_*m*_ = 1 for b- and y-ions and *w*_*m*_ = 0.2 for neutral losses. We consider neutral losses of carbon monoxide (CO, also known as “a-ions”), ammonia (NH_3_) and water (H_2_O) groups. If the precursor charge assigned to the spectrum is *>* 2, then fragments with charge *>* 1 are possible. In this case all peaks are replicated into lower-mass bins with the appropriate m/z. In particular, for a precursor of charge *n*, fragment ions up to charge *n* − 1 are considered. In most implementations of XCorr, if two or more predicted peaks fall in the same mass bin, then the intensity in that bin is the maximum of those peaks’ intensities. However, in order to facilitate calibration via dynamic programming, we wish to make the XCorr score function fully additive. Consequently, Equation 2 uses the sum of the peak intensities rather than the maximum (7).

We then predict a very simple theoretical spectrum from the sequence of the peptide. For this step, we use a discrete mass vector *B*, again using a bin width of 1.0005079 Da. For each possible backbone fragmentation of the peptide, *B* is populated with a single binary marker at the mass of the corresponding b ion.

The final XCorr score is a simple dot product between the evidence vector *E* and the theoretical spectrum *B*. Thus, the score is essentially the sum of the evidence for all the cleavage events across the length of the peptide.

### 2.2 The residue evidence score

The computation of the residue evidence score proceeds in the same steps outlined above for XCorr: pre-processing of the observed spectrum, aggregation of evidence, generation of a theoretical spectrum, and calculation of a score based on the evidence and the theoretical spectrum. Unlike the XCorr processing, which aggregates evidence into a vector indexed by m/z, the res-ev score aggregates evidence in a matrix indexed by m/z and amino acid.

1. The residue evidence score preprocessing employs a subset of the steps previously described for XCorr.
2. The intensity of each peak is replaced by its square root.
3. Peaks with intensity less than 5.0% of the maximum intensity peak are eliminated from the spectrum. Peaks within a 1.5 Th window of the precursor peak (assuming’remove-precursor-peak=T’) are also eliminated.
4. The spectrum is divided into 10 equal length segments, and the intensities within each segment is normalized so the maximum intensity in the segment is 50.

There are two important changes in this preprocessing compared to that for the XCorr score function. First, discretization of the peaks’ mass values is deferred until after the high-resolution residue evidence has been quantified and aggregated. Second, the final subtractive step, which induces a cross-correlation penalty in the XCorr score function, is omitted altogether. Another difference is the addition of two peaks, representing the *m/z* of the N-terminal group and the *m/z* of the precursor minus the C-terminal group (typically a hydroxyl group). Both of these peaks have intensities of zero.

Next, we quantify and aggregate evidence for each type of residue inducing a b-ion cleavage at each possible m/z bin. The residue evidence is defined as follows. Let an arbitrary pair of MS2 peaks *A* and *B* have measured masses *m*_*A*_ and *m*_*B*_, such that *m*_*A*_ > *m*_*B*_ and the difference in mass *m*_*diff*_ = *m_A_ − m_B_*. We say evidence *exists* for a (charge 1+) b ion fragment with mass *m*_*A*_, terminating in amino acid residue *X*, if

the deviation abs(*m_diff_ − m_X_*) *< m_tol_*, where *m*_*tol*_ is the maximum deviation tolerated between *m*_*diff*_ and *m*_*X*_. In practice, *m*_*tol*_ is on the order of the mass spectrometer’s MS2 resolution. The *magnitude r* assigned to this residue evidence is scaled so as to reward small deviations:

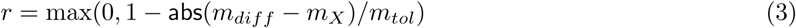

i.e., *r* takes a value of 1 when the deviation is 0, and 0 when the deviation is equal to or greater than *m*_*tol*_. This magnitude *r* is then multiplied by the sum of the rank intensities of the two peaks, prior to being stored.

Residue evidence is stored in a two-dimensional *residue evidence matrix R*. The columns of *R* are indexed by discretized masses *m*_*j*_, and the rows or *R* correspond to the amino acids *a*_*i*_ found in the peptide database (typically around 20 rows). The increment of evidence *r* generated according to Eq. 3 is added to element *R*_*aimj*_, where *a*_*i*_ = *X* and *m*_*j*_ is obtained by discretizing *m*_*A*_ with a bin width *W* = 1.0005079 Da.

Each pair of peaks *A* and *B* is also considered as a putative pair of charge 1+ y ions, charge 2+ b ions, and charge 2+ y ions, and additional residue evidence is generated using appropriate modifications to Eq. 3. In all cases, however, the evidence is added to the element of *R* indexed by the discretized mass of the corresponding charge 1+ b ion, so that all evidence related to a particular locus of fragmentation is aggregated together.

Once all possible residue evidence has been accumulated into matrix *R*, the values in *R* undergo a linear discretization to integer values, such that the minimum value in *R* is 0 and the maximum is some specified integer *r*_*max*_. This integer discretization ensures that scores will have integer values, which is required for the subsequent dynamic programming.

The theoretical spectrum corresponding to a candidate peptide is very simple. For each possible prefix sequence of the peptide a tuple is created, consisting of two elements: the identity of the prefix’s C-terminal amino acid and an integer formed by discretizing the prefix mass with bin width *W* = 1.0005079. For a candidate peptide *P* of length *n*, the full representation *B* then consists of *n* − 1 such tuples: {(*a_k_, m_k_*)} _*k*=1*…n−*1_.

Finally, assume that candidate peptide *P* of length *n* has a minimal binary representation *B* as described

above. Then the residue evidence score Ψ between *P* and spectrum *S* is the sum of elements selected from the residue evidence matrix *R* (derived from *S*) according to the tuples in *B*:

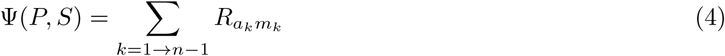

### 2.3 Calibrating the residue evidence score via dynamic programming

The following assumes a spectrum *S* with precursor mass is *m*_*S*_ is being scored.

Let *P* ^(1*→n*)^ be a peptide of length *n*, with mass *m*^(1*→n*)^ = *m*_*S*_ and amino acid sequence *a*_1_*, a*_2_*, …, a_n_*. Because Ψ(*P, S*) is additive, the score for matching *S* with *P* ^(1*→n*)^ can be obtained by first calculating the score for the prefix sequence *P* ^(1*→n−*1)^ = *a*_1_*, a*_2_*, …, a_n−_*_1_, then adding the evidence *r* = *R*_*anms*_ from the residue evidence matrix *R*. Note that this process is equally valid for any subsequence *P* ^(1*→k*)^ = *a*_1_*, a*_2_, …, *a*_*k*_ with mass *m*^(1*→k*)^,

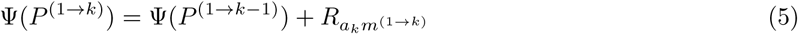

Let *C*_*s,m*_ be the count of peptides with mass *m* that produce a discretized score *s*. If (hypothetically) all the peptides have the same terminal amino acid *a* with mass *m*_*a*_, then we would have

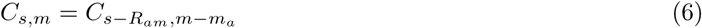

Allowing for all naturally occurring amino acids *a_i_ ∈ A*, with masses *m*_*ai*_, the count becomes

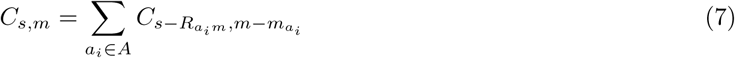

Since Ψ(*P, S*) is additive, Eq. 7 is valid for all masses 1 ≤ *m ≤m*_*S*_. Eq. 7 defines the basic recursion of the DP.

The DP computation of *C* is conducted in a two-dimensional array, where the rows are indexed by *s* and the columns by *m*. The number of rows is determined by an estimate of the largest possible score for *S*:

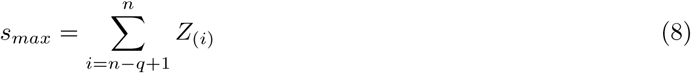

where *Z*_(*i*)_ refers to the sorted column maxima from *R* and 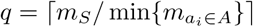.

We initially set:

- *C*_*0,T_N_*_← 1, where *T*_*N*_ is the mass of the N-terminal group.
- *C*_*s,m*_ ← 0 for all *s* ≠ 0 or *m* ≠ 1. This includes a range of indices *s <* 1 and *m <* 1 that are accessed during the DP.

The elements of the array are then computed sequentially:

~~~
**for** *m* = *T*_*N*_ to *m*_*S*_ − *T*_*C*_ **do**
        **for** *s* = *s*_0_ to *s*_*max*_ **do**
                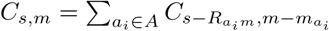
        **end for**
**end for**
~~~

Above, the values *T*_*N*_ and *T*_*C*_ represent the masses of the N-terminal and C-terminal groups, respectively. Hence, the last column of the matrix *C* typically represents mass *m_S_ −* 17, since the C-terminal group is usually a hydroxyl. This column holds the desired distribution of *X*_*R*_ over all possible peptides consistent with *m*_*S*_.

By using Eq. 7 in the DP, we make the assumption that all peptides are *a priori* equally likely. This is not biologically plausible, and, in fact, leads to distributions of Ψ(*P, S*) that lack appropriate statistical properties. This problem can be solved by considering the relative abundances of amino acids in the recursive counting:

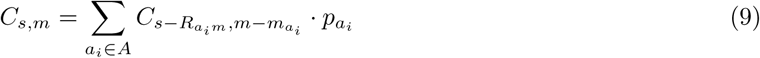

where *p*_*ai*_ is the probability of finding amino acid *a*_*i*_ in a large collection of naturally occurring peptides, with 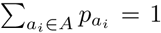.Note that it may be important to use different estimates of *p*_*ai*_ for the N-terminus, C-terminus, and non-terminal positions, depending on the specificity of the enzyme used for digestion.

Assume we have calculated, using DP, the distribution of scores *C*_*s,mS*_ over all possible peptides for spectrum *S*, where 0 ≤ *s ≤ s_max_*. Then the p-value relative to this distribution for a specific peptide *P*, matched to *S* with residue evidence score *ψ* = Ψ(*P, S*), is

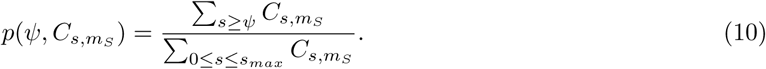

These p-values can be used in place of raw residue evidence scores during a standard database search.

### 2.4 Combining correlated p-values

The res-ev p-value and the XCorr p-value provide complementary yet not fully independent estimates of the quality of a given peptide-spectrum match (PSM). Accordingly, we employ a previously described method for assigning a p-value to the product of *n* correlated p-values (11), using the following equation:

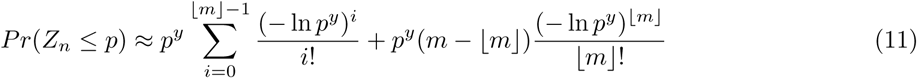

where *n* is the number of p-values being multiplied (in our case, *n* = 2), *Z*_*n*_ is the product of the p-values, *m* is a parameter that can range from 1 to *n*, and *y* = *m/n*. The value of *m* indicates the degree of correlation among the *n* p-values, where total correlation (i.e., identical p-values) corresponds to *m* = 1, and total independence corresponds to *m* = *n*. In this setting, we used decoy p-values to empirically estimate *m* = 1.2 (Supplementary Figure 1) by minimizing the previously described error function:

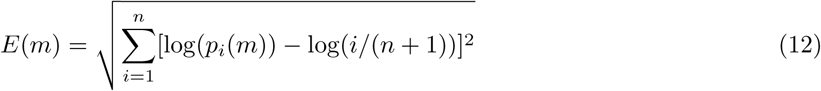

where *p*_*i*_(*m*) is the *i*th largest p-value (of the product of p-values) in a set of *n* decoy p-values. Intuitively, this error function attempts to minimize the difference between the observed p-value distribution and an ideal, uniform distribution.

### 2.5 Data sets

We used four previously described tandem mass spectrometry data sets to validate our methods (Table 1). The data sets were selected to represent a diversity of both sample types and instrument types.

**Table 1:**
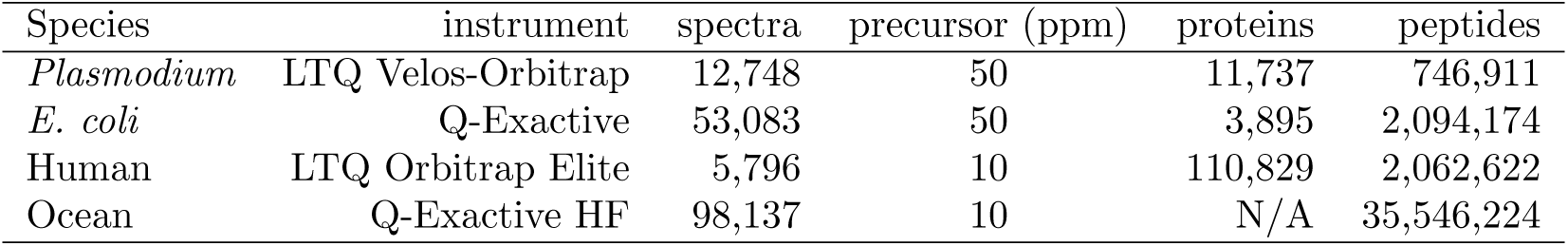
Mass spectometry datasets. The database used for the ocean data set is comprised of individual peptides derived from high-throughput sequencing reads, rather than full-length proteins.

#### Plasmodium falciparum fraction (13)

*P. falciparum* 3D7 was grown at 37^*◦*^ Celsius in RPMI-1640 culture medium. Following synchronization, infected cells were lysed using saponin. An 8 M urea lysis buffer was used to create parasite extracts, which were then reduced and alkylated. Proteins were digested using Lys-C, and the resulting peptides were labeled with TMT. Following TMT labeling, strong cation exchange chromatography (SCX) was use to fractionate the sample into 20 fractions. Fractions were analyzed on a LTQ Velos-Orbitrap mass spectrometer (Thermo Scientific). All MS1 and MS2 scans were acquired at high resolution. The data from fraction 13 was used in this study and contained 12,748 scans. The protein database used in the database search was downloaded from NCBI in October 2013 (*Plasmodium falciparum 3D7*).

#### Ocean metaproteome(14)

Water samples from the northern Chukchi Sea bottom waters were collected in the summer of 2013. To remove larger eukaryotes, each 15 L water sample was prefiltered through a 10 *µ*m and then a 1 *µ*m filter. The remaining liquid was collected onto a glass fiber filter and frozen. Cells were lysed using bead beating in 6 M urea. A total of 100 *µ*g of total protein were used for digestion. Prior to digestion, 300 *η*g of human ApoA1 protein was added and then the sample reduced and alkylated. Proteins were digested with trpysin and then desalted. Peptide separation was conducted using a NanoAquity HPLC with a 4 cm precolumn and a 30 cm analytical column. Peptides were eluted at a rate of 300 *η*L/min for 2 hours using a nonlinear gradient. Data was collected on a Q-Exactive HF (Thermo Scientific). The mass spectrometer was operated in a Top 20 data-dependent acquisition mode with a 5 second dynamic exclusion window. Ions only between 400-1600 *m/z* were collected. This resulted in a dataset with 98,137 scans. The database used in the database search consists of a metapeptide database that was derived from shotgun metagenomic sequencing of the same ocean sample (https://noble.gs.washington.edu/proj/metapeptide/metapeptidesCS.fasta). Briefly, a metapeptide database is a peptide database whose sequences are derived from raw read sequences that have been translated into peptides in all six reading frames (14).

#### Human fraction (15)

Histologically normal adrenal gland tissue from three deceased individuals were pooled together using equal amonts of protein from each donor. Samples were lysed using SDS. The protein sample was fractionated by SDS-polyacrylamide gel electrophoresis (SDS-PAGE). Then the protein bands were destained, reduced, and alkylated. Protein digestion was peformed using an in-gel trypsin digestion. Following digestion, the peptides were desalted. The resulting peptide sample was separated with a 60 min linear gradient using reversed-phase liquid chromatography on an Easy-nLC II nanoflow liquid chromatography system (Thermo Scientific). Data was acquired using a LTQ-Orbitrap Elite (Thermo Scientific). The mass spectrometer was operated in a Top 20 data-dependent acquisition mode with a 30 second dynamic exclusion window. MS1 scans were acquired at a mass resolution of 120,000 at 400 *m/z*. MS2 scans were acquired at a mass resolution of 30,000 at 400 *m/z*. This study used data from the first fraction, which contained 5,796 scans. Each chromosome’s “protein.faa” was downloaded from RefSeq (ftp://ftp.ncbi.nlm.nih.gov/refseq/Hsapiens/mRNAProt) in September 2016 and concatenated to form the human protein database.

#### E. coli fraction(16)

A yfgM knockout strain and WT strain of *E. coli* MC4100 was cultured at 37^*◦*^ Celsius in LB broth (Difco, Sparks, MD). Cells were harvested using centrifugation once the OD_*600nm*_ reached ~.08. Cellular pellets were suspended in a lysis buffer and then lysed by rapidly passaging the cells through a hypodermic syringe needle and by sonication. Proteins were then reduced, alkylated, and then reduced a second time. A 4 hour digestion step with Lys-C was followed by an overnight trypsin digestion. The resulting peptides were chemically labeled using stable isotope dimethyl labeling. The yfgM knockout lysate was labeled with the “Medium” isotope and the the WT sample was labeled with the “Heavy” isotope. These lysates were mixed together in a 1:1 ratio and then fractionated into 45 fractions using strong cation exchange. Samples were analyzed on a Q-Exactive (Thermo Scientific) coupled to a Easy UHPLC (Thermo Scientific) system. Peptides were eluted during a 3 hour gradient with a flow rate of 100 *η*L/min. The Q-Exactive instrument was operated in a Top 20 data-dependent acquisition mode. MS1 scans were acquired at a mass resolution of 35,000. MS2 scans were acquired at a mass resolution of 17,500. The 21^st^ fraction was used for this study and contained 53,083 scans. The protein database used in the database search was downloaded from Uniprot in December 2017 (*Escherichia coli* str. K-12 substr. MC4100).

### 2.6 Target-decoy evaluation

For this work, we used the following publicly available database search engines.

- Crux version 3.1 (http://crux.ms; linux version) was used to generate combined p-value, res-ev p-value, XCorr p-value, and high-resolution XCorr (7, 17, 18)
- MS-GF+ version v2016.12.12 (https://omics.pnl.gov/software/ms-gf) (19)
- MS Amanda version 1.0.0.7504 (http://ms.imp.ac.at/?goto=msamanda; linux version) (20)
- Morpheus version 272 (http://cwenger.github.io/Morpheus/; linux version) (4)

We took great care to ensure a fair comparison of results across all database search engines. One of the more important ways we accomplished this goal was to try to guarantee that all the database search engines considered a common set of target and decoy peptides. To this end, we took several non-standard steps in our analysis. First, we predigested our protein fasta files *in silico* using the tide-index tool in Crux. This predigestion did not include suppression of cleavage by proline, because not all search engines use this rule. Decoy peptides were generated by tide-index by shuffling the amino acid sequence of each peptide, leaving the N-terminal and C-terminal amino acids in place. For this digestion, no missed cleavages were allowed, and N-terminal peptides with a leading methionine were included in two copies, with and without the methionine. Peptides shorter than six amino acids and peptides with one or more non-enzymatic termini were not considered. The resulting target and decoy peptides were placed into a new.fasta file. The words ‘target’ and ‘decoy’ were appended to the peptide headers of the peptides in their respective .fasta files. Then the target and decoy .fasta file were concatenated to create a target-decoy database for MS Amanda.

Because it is not possible to turn off N-terminal methionine clipping in MS Amanda, in order to ensure that the other three database search tools were exposed to the same peptides as MS Amanda, we then subjected the predigested .fasta files to a second round of “digestion”. In this second round, no missed cleavages were allowed, N-terminal methionines were allowed to be clipped, and peptides shorter than five amino acids were removed. Since the .fasta files that went through the second round were pre-digested, this next step only performed clipping of N-terminal methionines. The resulting target and decoy peptide files were then concatenated to create a target-decoy database for MS-GF+, Morpheus, and Crux. This predigestion strategy ensured that all search engines considered the same set of candidates. Note that, subsequent to the searches, we also checked that each detected peptide was indeed present in the .fasta database.

In addition to ensuring the database search engines considered the same set of targets and decoys, we tried to match the experimental parameters in each database search as exactly as possible. We removed any MS2 scans that had fewer than 10 peaks in it. All searches were run with full digestion (*i.e.* no missed cleavages). No non-enzymatic termini and no isotope errors were allowed. The maximum precursor charge was set to 25 (*E. coli*), seven (human and ocean), or nine (*Plasmodium*). The *E. coli*, human, and ocean sample were run with trypsin as the digestion enzyme, while the *Plasmodium* sample was run with Lys-C as the digestion enzyme. The proline rule was ignored for all runs. The precursor mass tolerance was set to 50 ppm for the *E. coli* and *Plasmodium* runs and 10 ppm for the human and ocean runs. We set the fragment mass tolerance at 0.02 Da for combined p-value, res-ev p-value, MS Amanda, and Morpheus for all four datasets; however, we were unable to set the fragment mass tolerance for MS-GF+ as it is not a user-level parameter. For MSGF+, we can somewhat control the fragment mass tolerance by correctly setting the user-level parameters of ‘inst’ and ‘m’. For the human and *Plasmodium* dataset, we set ‘inst’ to 1 and ‘m’ to 3. These settings correspond to high-resolution MS2 scans that were generated by HCD. For the ocean and *E. coli* sample, we set ‘inst’ to 3 and ‘m’ to 3. These settings correspond to high-resolution MS2 scans that were generated by a Q-Exactive. For all four datasets, we allowed a fixed carbamidomethyl modification to cysteine and a variable methionine oxidation modification. In addition we allowed a variable light, intermediate, and heavy dimethyl label (28.0313, 32.0564 and 36.0757 Da, respectively) on lysines and the N-terminus for the *E. coli* run. For the *Plasmodium* run, we included a fixed TMT modification (229.16293 Da) on lysines and the N-terminus. Methionine clipping was turned off for Crux, Morpheus, and MS-GF+ since the input .fasta file already contained clipped peptides. All search engines runs were done in target-only mode since the peptide headers in the concatenated target-decoy .fasta file already denoted whether it was a target or decoy peptide. The exact commands used to run the database searches can be found in Supplementary File 5.

A custom R script (Supplemental File 3) was used to combine the results of the various search engines together into a single table. Each row represents the PSM that each database search detected for a particular scan. For each row (scan) the combined p-value, res-ev p-value, XCorr p-value, SpecEvalue (MS-GF+), weighted probability (MS Amanda), and Morpheus scores are listed in Supplemental File 2. In addition, the peptide that each score function detected, and whether that peptide is a target or decoy, is also listed. The value ‘NA’ is placed into empty cells that result from one score function scoring a scan and another score function not scoring that particular scan. This phenomenon is due to each program having a different threshold for the minimum number of peaks required to score a scan. A second R script used the PSM table as input to calculate false discovery rates and generated the plots for this publication (Supplemental File 4). We used the following false discovery rate equation: FDR = (number of decoys + 1) / number of targets (21).

### 2.7 Percolator analysis

For the Percolator analysis, the Tide search was performed as described previously, except that during index creation, the “digestion” option was set to “partial-digest” and one missed cleavage was allowed (“missed-cleavages=1”). This setting allows Percolator to more effectively re-rank various types of PSMs, while taking into account their digestion conditions. We then applied the Crux implementation of Percolator directly to the Tide search results. The resulting feature vector contains, in addition to the standard Percolator features, three separate scores for each PSM: the negative logarithms of the combined p-value, res-ev p-value, and XCorr p-value. All default Percolator parameters were used except that ”only-psms” was set to true. Note that PSM-level FDR is estimated by Percolator using target-decoy competition (including the +1 correction to the number of decoys).

## 3 Results

### 3.1 Statistical validation of residue-evidence p-value

For any observed spectrum, we can use dynamic programming to determine the exact distribution of residue-evidence scores that result from each possible peptide sequences whose discretized mass matches the discretized precursor mass. We can then compare the score from a particular PSM to this distribution and calculate a p-value, i.e., we compute the probability of observing a residue-evidence score greater than or equal to the score of a particular PSM.

To test the validity of the resulting p-values, we searched real data against a decoy database. Specifically, we searched the *Plasmodium* data set against a shuffled *Plasmodium* database (see Methods for details) using a wide precursor mass tolerance of 3 Da. Because the decoy peptides have been shuffled, we expect all of the resulting PSMs to be incorrect. Hence, in this setting, our p-values should be uniformly distributed; i.e., the probability of observing a p-value less than or equal to, say, 0.05 should be 5%. A quantile-quantile plot (Figure 2A) of the calculated p-values against the rank of the p-values confirms that the residue-evidence p-values are generally uniform. However, we noticed a trend away from *y* = *x* among large p-values (horizontal line in the upper right hand corner of Figure 2A), as well as an overall upward shift of the p-value distribution shown in the figure. These two phenomena arise because of the discrete nature of the res-ev score. In practice, many PSMs result in a residue-evidence score of 0, leading to an inflation of p-values of 1.0 and a consequent decrease in the remaining p-values. Overall, the near uniformity of the empirical res-ev p-values indicates that they provide an accurate assessment of the statistical confidence associated with a given PSM.

**Figure 2:**
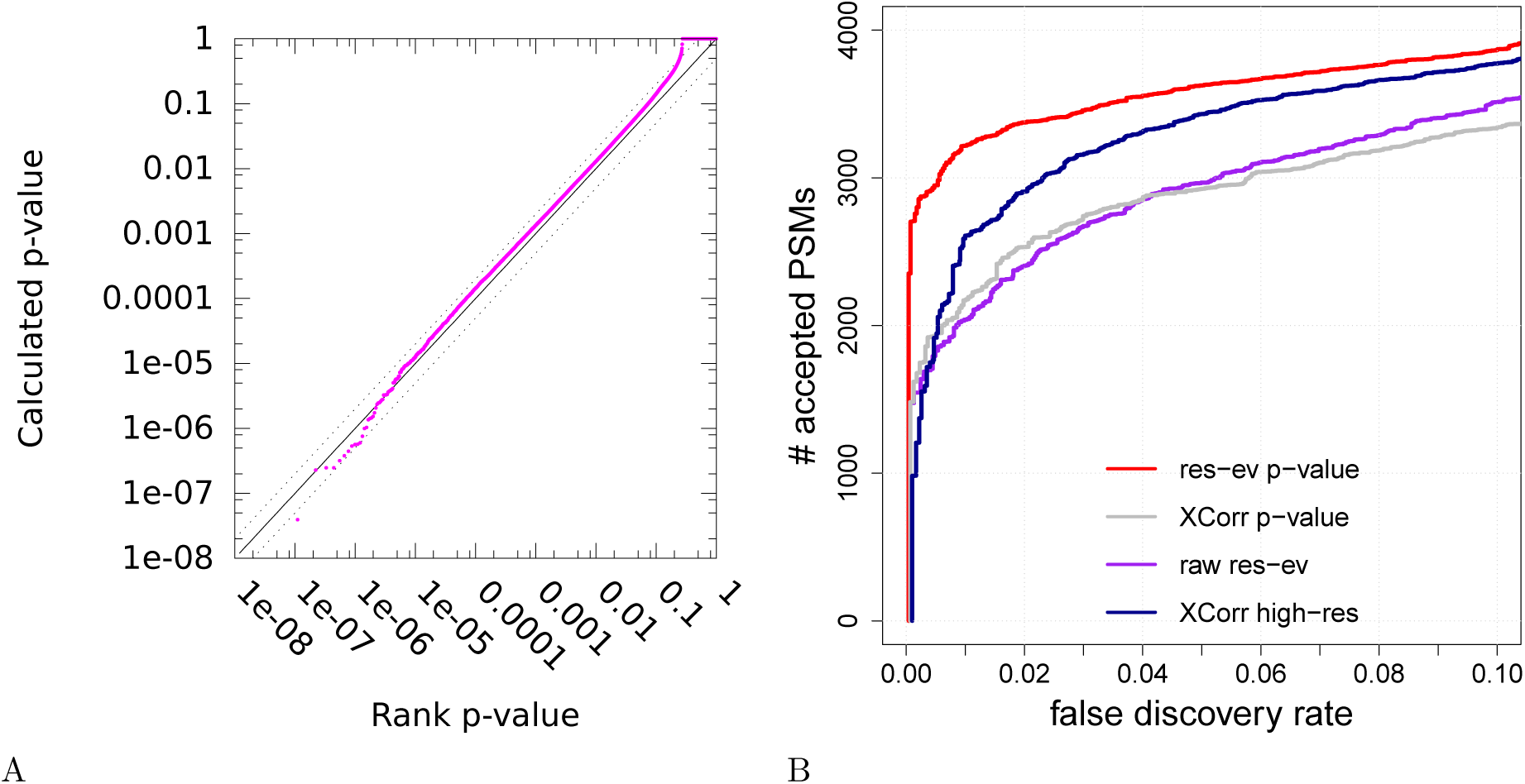
Calibration of the residue evidence score via dynamic programming. (A) The figure plots the p-value, as calculated via dynamic programming, versus the rank p-value, for decoy PSMs from a *Plasmodium* dataset. The lines *y* = *x* (solid line), *y* = 2*x* (dotted line) and *y* = 0.5*x* (dotted line) are included for reference. (B) The figure plots, for the *Plasmodium* dataset, the number of PSMs accepted as a function of *q*-value threshold, for four different database search methods: XCorr calibrated via dynamic programming using low-resolution m/z bins, uncalibrated XCorr using high-resolution m/z bins, uncalibrated res-ev, and res-ev calibrated via dynamic programming.

### 3.2 Residue-evidence works well for high-resolution data

Having established the validity of the res-ev p-value, we next sought to measure the statistical power of the score function in the context of a real database search. For this test, we again used the *Plasmodium* data set, but we searched against a concatenated database of both real (“target”) and shuffled (“decoy”) peptide sequences. From the resulting ranked list of PSMs, we used target-decoy analysis to estimate the false discovery rate (FDR) associated with each observed PSM score (22). We measured FDR out to a maximum of 10%, reasoning that higher FDR thresholds are not likely to be of much practical value. For comparison, we repeated our search with three other score functions: the uncalibrated res-ev score function, the uncalibrated high-res XCorr score function, and the XCorr p-value. Note that the latter necessarily discards the high resolution of the fragment m/z axis, because the XCorr dynamic programming procedure requires ~1 Da bins.

The results (Figure 2B) clearly show that the res-ev p-value score function outperforms the three competing methods. Focusing on the commonly used FDR threshold of 0.01, we see that the res-ev p-value detected 3,217 PSMs. This corresponds to an increase of 1178 (57.78%) PSMs relative to the raw residue-evidence score, 1,047 (48.25%) PSMs relative to the XCorr p-value and 609 (23.35%) PSMs relative to high-resolution XCorr. Thus, this experiment suggests that taking simultaneous advantage of statistical calibration and high-resolution data improves preformance.

A complementary measure of the quality of a PSM score function is the “target match percentage” (TMP), which is defined as the fraction of spectra for which the top-scoring match involves a target peptide (23). For a perfectly random score function, we expect the TMP to be ~50%. The best possible TMP is 100%; however, in practice any real data set will contain spectra that cannot be identified, either because the corresponding generating peptide is not in the given peptide database or because the spectrum was generated by a non-peptide contaminant. TMP provides a measure of the quality of a score function that is independent of a score function’s calibration. This is because the TMP never involves comparing scores for PSMs involving different spectra. Hence, the distribution of PSM scores for spectrum *A* can be dramatically different from the distribution of PSM scores for spectrum *B*, but the TMP achieved by the score function can still be high.

In the *Plasmodium* TMP analysis, the res-ev p-value (and, by definition, also the raw res-ev score) achieved the best TMP of 65.08% (8,172 PSMs). High-resolution XCorr yielded the second best TMP of 64.20% (8,061 PSMs). Not surprisingly, XCorr p-value, which discards high-resolution m/z information, had the worst TMP (63.94%, or 8,029 PSMs).

In order to better understand the differences in scoring between XCorr and residue-evidence, we looked at several spectra where XCorr p-value and res-ev p-value greatly disagreed on the significance of their best PSMs. Figure 3 shows two such spectra (scans 5468 and 11156) from the *Plasmodium* dataset that have been annotated by both XCorr and residue-evidence.

**Figure 3:**
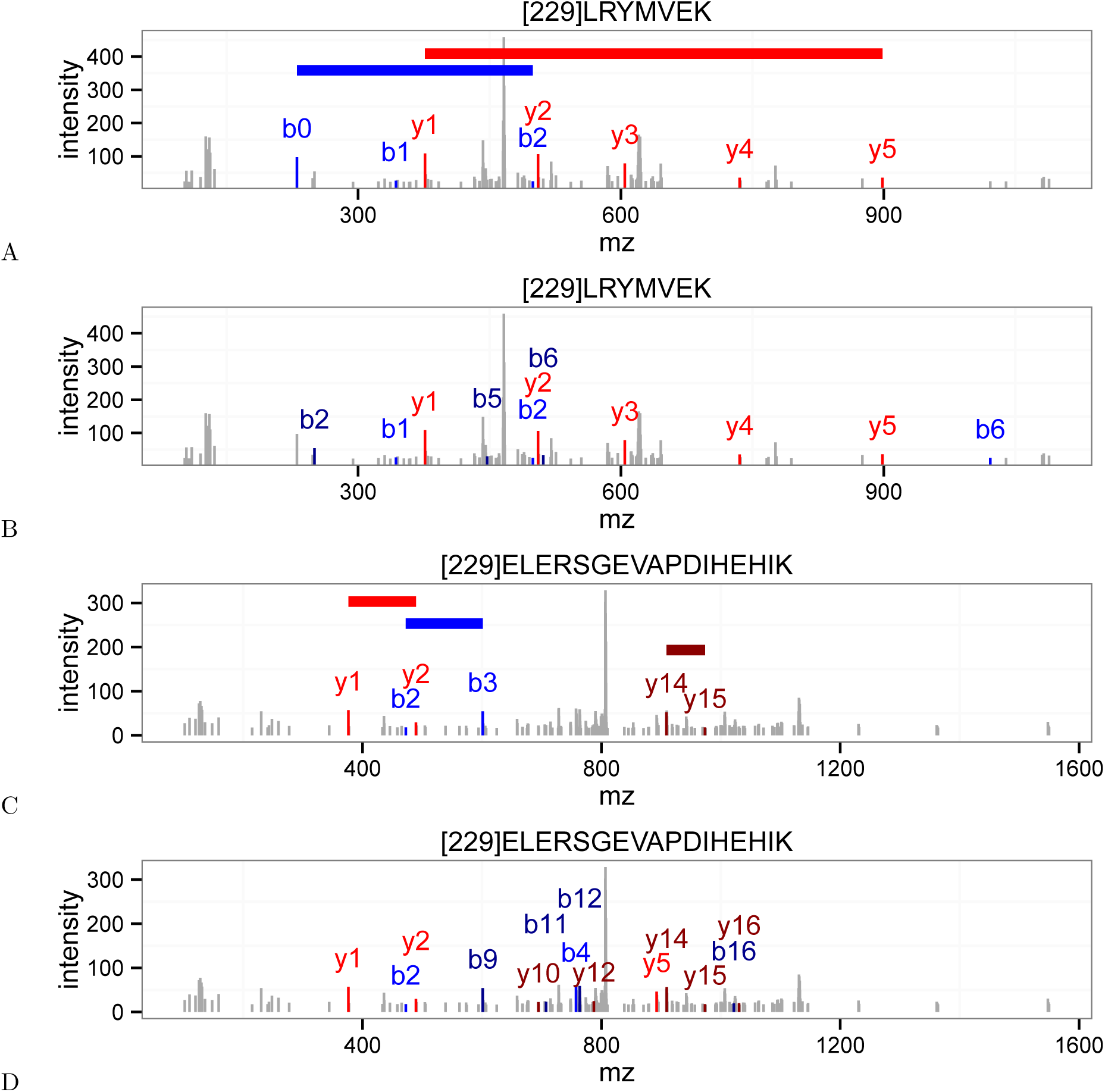
Disagreements between the XCorr score and the residue-evidence score. (A) An annotated *Plasmodium* spectrum (scan 5468) that received a low (i.e., good) p-value from the residue-evidence score. Colored horizontal lines indicate the locations of peak-pairs that contribute to the residue-evidence score. (B) Same as (A), but annotated using XCorr. This scan received a high XCorr p-value. (C) *Plasmodium* scan 11156, annotated with res-ev, with a high p-value. (D) Same as (C), but annotated using XCorr, with a low p-value. In each panel, peaks colored in blue, dark blue, red, and dark red represent b+1, b+2, y+1, y+2 ions, respectively.

Scan 5468 (Figure 3A–B) corresponds to a case where the PSM is given a small residue-evidence p-value and a large XCorr p-value. Specifically, although this scan was assigned the same peptide (LRYMVEK) by both score functions, the resulting PSM received a p-value of 5.13 × 10^*−*4^ (0.32% FDR) from residue-evidence and a p-value of 7.20 × 10^*−*1^ (48.50% FDR) from XCorr. The source of this difference is not immediately obvious, because the numbers of peaks annotated by the two score functions are similar. The only additional ions that XCorr identifies over residue-evidence are the doubly charged b2, b5, and b6 ions and the singly charged b6 ion. This PSM scores well according to the res-ev score function because the spectrum contains two long “ladders” of consecutive peaks (y1 to y5, and b0 to b2). These ladders are particularly unlikely according to a null model in which each peak is treated independently. Conversely, the poor score from XCorr may arise because most of the annotated peaks have low intensities. Note that the res-ev score did not annotate the doubly charged b5 and b6 ions because the mass difference between these two peaks was too different from the mass of glutamate. This speaks to the power of using high resolution on the MS2 m/z axis.

In contrast, scan 11156 (Figure 3C–D) illustrates why some PSMs score well using XCorr and poorly using residue-evidence. This scan was assigned the same peptide (ELERSGEVAPDIHEHIK) by both score functions, but the resulting PSM received a high p-value (8.45 × 10^*−*1^, 47.4% FDR) from residue-evidence and a low p-value (1.11 × 10^*−*2^, 3.4% FDR) from XCorr. The disparity between the two score functions arises because, using residue-evidence, only three pairs of peaks contribute to the score. In contrast, the XCorr score includes individual components corresponding to sixteen different fragment ions (b2, b3, b4, b7, b9, b11, b12, b16, y1, y2, y5, y10, y12, y14, y15, and y16). In Figure 3D, there are only fourteen colored fragment ions because the b3 and b9 ion correspond to the same peak and the b7 and the y16 ion also correspond to the same peak. The lack of a long “ladder” of successive b- or y-ion peaks keeps the residue-evidence score low, relative to XCorr.

### 3.3 Combining the two scores yields improved power

The two scans in Figure 3 clearly suggest the need for a score function that can combine the res-ev p-value and the XCorr p-value, thereby potentially correctly identifying both scans 3667 and 11156. Estimating the p-value for the product of a pair of independent p-values is relatively straightforward; however, in our case, the res-ev and XCorr p-values are clearly not independent since they are derived from the same PSM. To verify this lack of independence, we computed the res-ev p-value and XCorr p-value for four different data sets. These data sets were selected for diversity: they represent different proteome sizes, digestion enzymes (trypsin and Lys-C), instruments (LTQ Orbitrap Velos, Q-Exactive, and LTQ-Orbitrap Elite), and instrument resolutions. The strong *y* = *x* component in each of the resulting density plots (Figure 4, top row) indicates a strong lack of independence. We also note that, in general, res-ev shows an enrichment of very small p-values compared to XCorr.

**Figure 4:**
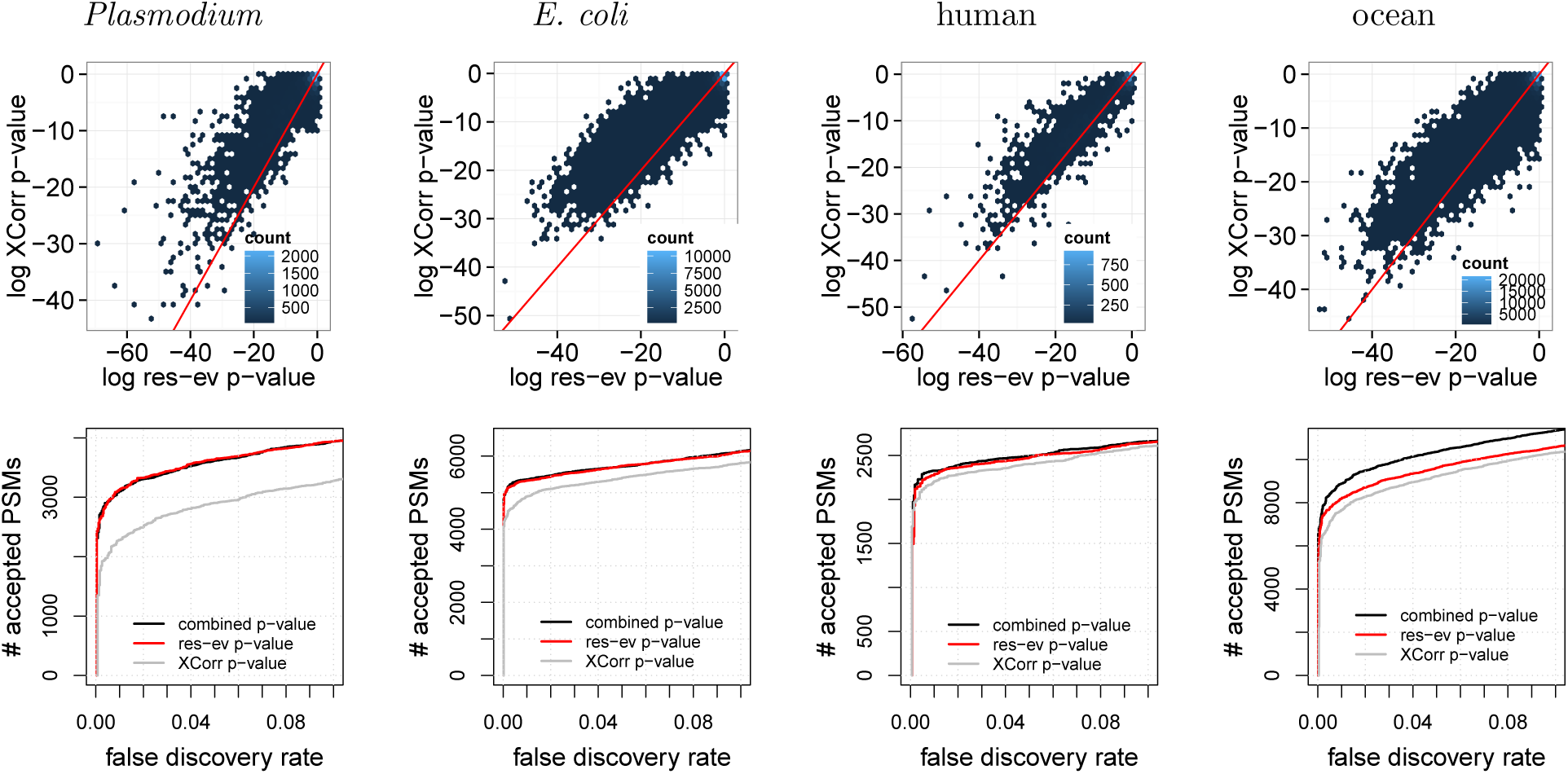
Combining XCorr and res-ev. (Top row) Each panel plots, for a specified dataset, a density plot of p-values from XCorr (y-axis) versus res-ev (x-axis). The points are binned into hexagons, and the color of each hexagon represents the number of points within each bin. The red line represents *y* = *x*. (Bottom row) Each panel plots the number of PSMs accepted as a function of FDR threshold, for three different database search methods: XCorr p-value, res-ev p-value, and combined p-value.

To combine these two scores, we applied a previously described method for estimating the statistical significance of the product of correlated p-values (24) (see Methods for details). We hypothesized that the resulting combined p-value would perform better than res-ev or XCorr because these two score functions take advantage of different types of evidence in the spectrum: the res-ev p-value focuses on adjacent pairs of peaks, whereas the XCorr p-value focuses on single peaks.

To test this hypothesis, we compared the performance of res-ev p-value, XCorr p-value and the combined p-value on four data sets. The results (Figure 4, bottom row) show that the combined p-value is generally the best-performing method. Notably, however, in three out of the four cases, the performance of combined p-value is comparable to res-ev p-value. In two out of the three cases, combined p-value identified only 35 (0.67%, *E. coli*) and 51 (2.24%, human) more PSMs than res-ev p-value at a 1% FDR threshold. In the third case, combined p-value did marginally worse than res-ev p-value: at a 1% FDR, combined p-value detected 29 (0.94%) fewer PSMs than res-ev p-value for the *Plasmodium* dataset. Only in the ocean dataset did the combined p-value yield a large increase in performance (817 PSMs, or 11.01%). Conversely, XCorr p-value tends to perform poorly in all cases. Overall, these results suggest that combining res-ev with XCorr does not lead to decreased performance and occasionally yields a performance increase relative to using the res-ev p-value alone.

**Table 2:**
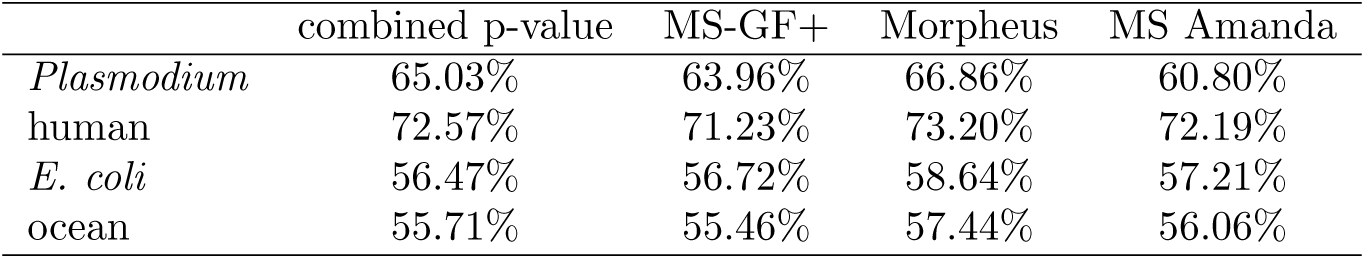
Target match percentages. The TMPs of four score functions (rows) for four datasets (columns). The TMP is defined as the percentage of spectra that match a target peptide.

|  | combined p-value | MS-GF+ | Morpheus | MS Amanda |
| --- | --- | --- | --- | --- |
| *Plasmodium* | 65.03% | 63.96% | 66.86% | 60.80% |
| human | 72.57% | 71.23% | 73.20% | 72.19% |
| *E. coli* | 56.47% | 56.72% | 58.64% | 57.21% |
| ocean | 55.71% | 55.46% | 57.44% | 56.06% |

### 3.4 Comparison with existing methods

Finally, we compared the combined p-value with three existing methods that take advantage of high-resolution tandem mass spectra: MS Amanda, Morpheus, and MS-GF+. MS Amanda and Morpheus are designed to take advantage of high-resolution tandem mass sectra but are not statistically calibrated. MS-GF+, like res-ev p-value and combined p-value, takes simultaneous advantage of statistical calibration and high-resolution MS2.

We found that, in general, combined p-value and MS-GF+ outperformed MS Amanda and Morpheus over the entire 0–10% FDR range (Figure 5A). For example, combined p-value detected 1781 (135.33%), 711 (44.02%), and 2414 (37.33%) more PSMs than MS Amanda at a 1% FDR for the *Plasmodium*, human, and ocean samples, respectively. Similar respective improvements of 1479 (91.41%), 404 (21.02%), and 3074 (52.95%) were observed for the combined p-value relative to Morpheus, as well as for MS-GF+ relative to MS Amanda and Morpheus. The one exception was the *E. coli* dataset, where MS Amanda performed comparably to MS-GF+ and slightly better than the combined p-value. Nonetheless, we interpret the performance improvement that the combined p-value and MS-GF+ offer over MS Amanda and Morpheus as evidence of the value of having a statistically calibrated score.

**Figure 5:**
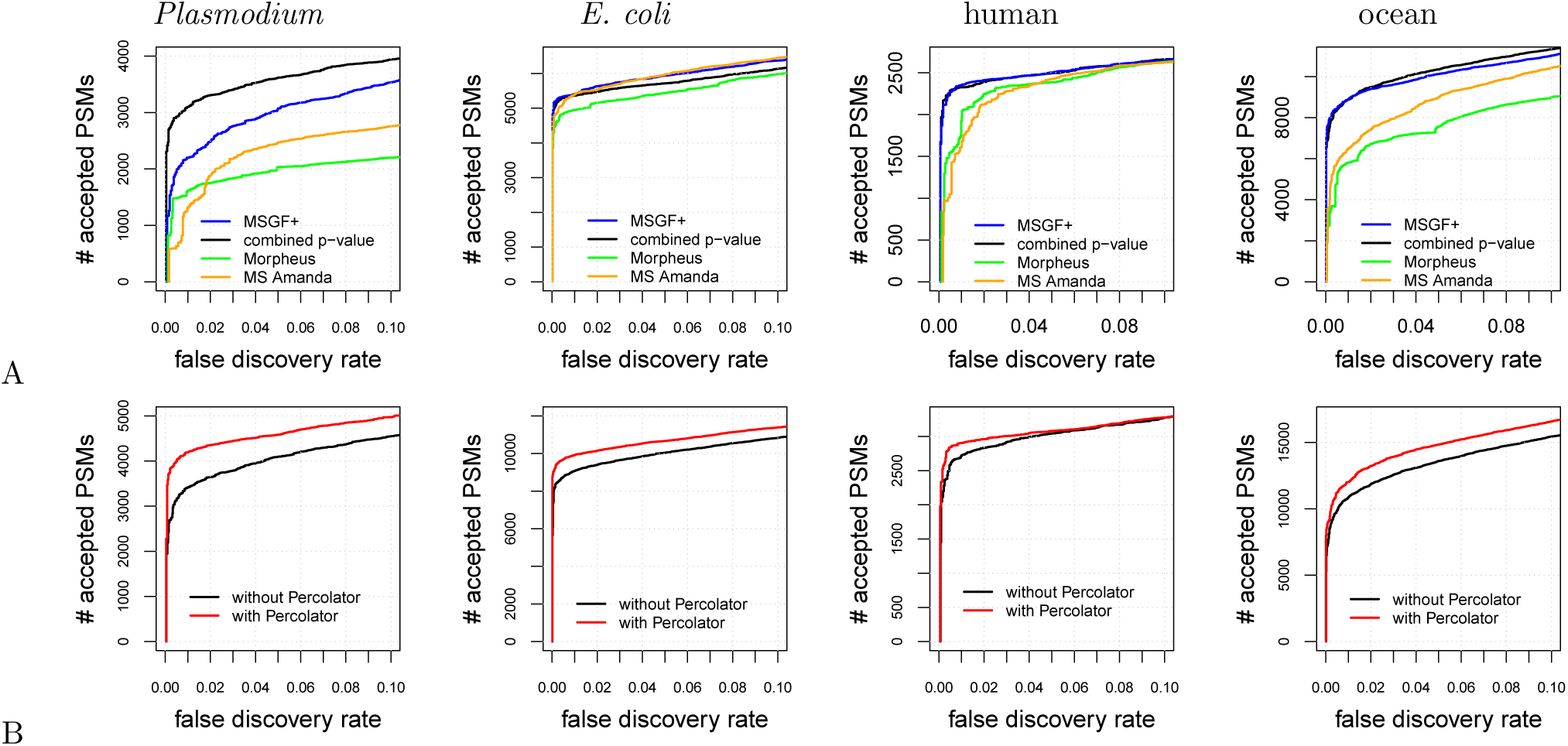
Comparison with existing methods. (A) The panel plots, for four datasets, the number of PSMs accepted as a function of FDR threshold for four different database search methods: MS-GF+, combined p-value, MS-Amanda, and Morpheus. (B) Similar to (A), but the two series correspond to the combined p-value with and without post-processing via Percolator.

When we focus on the comparison between the combined p-value and MS-GF+, no clear winner emerges. For three datasets, *E. coli*, human, and ocean, MS-GF+ marginally outperformed combined p-value at a 1% FDR threshold. Qualitatively, we observe on the *E. coli* dataset that the difference between MS-GF+ and the combined p-value is relatively small for low FDR thresholds—e.g., at a 1% FDR, MS-GF+ yields only 42 (0.78%) more PSMs than the combined p-value—but increases for larger FDR thresholds. Conversely, for the human dataset the difference between MS-GF+ and the combined p-value is largest (24 PSMs or 1.03%) at around 1% FDR, but then this difference all but disappears for large FDR thresholds. In contrast to the prior two datasets, for the ocean dataset MS-GF+ identfied 8 (0.09%) more PSMs relative to combined p-value at a 1% FDR threshold. However, at larger FDR thresholds, combined p-value consistently performed better then MS-GF+. Finally, for the *Plasmodium* dataset, we observed that combined p-value performed dramatically better than MS-GF+ across the entire FDR range, with an improvement of 899 PSMs (40.90%) at a 1% FDR.

Comparing the target match percentages of combined p-value, MS-GF+, Morpheus, and MS Amanda yielded unexpected results. In contrast to the comparison shown in Figure 5, where Morpheus consistently performed worse than the other score functions, in the TMP comparison Morpheus performed better than the other score functions. Morpheus’s TMP was higher than the second-best TMP by 1.83%, 0.63%, 1.43%, and 1.38% for the *Plasmodium*, human, *E. coli*, and ocean datasets, respectively. Among the remaining three methods, the TMP values for most of the data sets were remarkably similar to each other, spanning ranges of 71.23–72.57% (human), 56.47–57.21% (*E. coli*), and 55.46–56.06% (ocean), respectively. The TMP values were slightly more variable for the *Plasmodium* dataset, but even in this case, Morpheus was a clear winner. These results suggest that Morpheus is doing a very good job of identifying the correct candidate peptide for each spectrum, and perhaps suffers in the calibration of its scores from one spectrum to the next.

### 3.5 Using Percolator in conjunction with combined p-value improves power

In practice, in most proteomics experiments a post-processor such as Percolator (25) or PeptideProphet (26) is used to reanalyze the database search results to improve performance. Therefore, we tested whether the performance of combined p-value can be improved by post-processing via Percolator. We ran the combined p-value database search for all four datasets, as previously described, except that we allowed the peptide database to contain semi-tryptic peptides and peptides with one missed cleavage. This approach provides Percolator the opportunity to re-rank PSMs while taking into account their digestion conditions. Following the database search, we reanalyzed the database results with Percolator. We found that combined p-value with Percolator performed better than combined p-value by itself for all four datasets over the entire 0–10% FDR range (Figure 5B). At a 1% FDR, Percolator identifies an additional 778 (22.8%), 844 (9.30%), 182 (6.69%), and 1180 (10.89%) PSMs for the *Plasmodium*, *E. coli*, human, and ocean datasets, respectively.

## 4 Discussion

The residue-evidence p-value is reminiscent of the original XCorr score employed by SEQUEST but is designed to take simultaneous advantage of statistical calibration and high-resolution tandem mass spectra. By combining residue-evidence p-values with XCorr p-values, we obtain state-of-the-art performance in identifying tandem mass spectra. The resulting search engine is freely available in the open source Crux mass spectrometry toolkit (http://crux.ms).

The number of search engines available to process shotgun proteomics mass spectrometry data is large and growing (reviewed in (3)). Given this diversity of approaches and the different results produced by each search engine, it is only natural to attempt to search the same data with multiple methods and combine the results in a post-processing stage. Indeed, a wide variety of methods have been developed that adopt this approach, including methods that aggregate PSMs and then re-estimate FDRs on the aggregated results (27–29), combine statistical confidence measures (30), compute probabilities for each method and then combine these probabilities (31–34), or run a machine learning post-processor on the combined results (35). Our empirical results suggest that the res-ev score function and its combined p-value provide yet another complementary view of peptide-spectrum matching, which will likely add value in the context of such aggregation schemes.

One potential explanation for the relatively poor performance of MS-GF+ on the *Plasmodium* dataset is due to the unusual nature of the data itself. MS-GF+ uses a supervised machine learning algorithm to learn a scoring model. To perform well on the *Plasmodium* data might require a model that was trained on data with TMT labeling and digestion with Lys-C. In contrast, the combined p-value approach is invariant to properties of the data, such as digestion and labeling schemes.

A potential area for future work lies in the method for combining XCorr and res-ev p-values. We empirically set the parameter *m*, which represents the degree of dependency between the two types of p-values, to be 1.2. However, in principle this value could be re-estimated for each new data set, using a strategy similar to that shown in Supplementary Figure 1. However, our empirical results (not shown) suggest that the behavior of the method is not strongly dependent upon the choice of *m*.

The strong performance of the Morpheus search engine as measured by TMP suggests that this score does a good job of identifying the generating peptide for a given spectrum. On the other hand, the poor overall performance of Morpheus suggests that the score function is poorly calibrated. This is not surprising, because the score is simply the sum of two terms: the number of matched product ions, and the fraction of the observed peak intensities that can be assigned to matched products. Thus, longer peptides or spectra with more peaks will tend to achieve higher Morpheus scores on average. Our results suggest that a calibrated version of this score function should be able to achieve very good empirical performance.

In addition to proposing a new score function, we have provided a new benchmark for use in evaluating novel score functions. On the surface, it seems deceptively easy to compare results across score functions: run the score functions on the set same of input spectra and peptides and then compare the results. However, in reality, it is much harder to fairly compare score functions because search engines differ in many ancillary ways: digestion rules, decoy generation, etc. By merging all of the PSMs from different search engines, for a given dataset, into a single table indexed by scan number (Supplementary Tables 1-4), and by ensuring that all the reported peptides appear in a shared peptide list, we ensure that the performance evaluation focuses on properties of the score function, rather than less interesting properties of the digestion rules or candidate peptide selection procedure.

## Supplemental files

The following supplemental files can be found in PRIDE under accession PXD009265.

1. ***E. coli* peptide database (ecoliPSMDb.txt.gz).**The tab-delimited file contains the following columns: peptide sequence, peptide mass, an indication of whether the peptide is a target or decoy, and a comma-separated list of the IDs of proteins containing this peptide. Dynamic modifications are indicated using bracket notation. Static modifications are not indicated. Decoy peptides do not have protein IDs associated with them.
2. **Human peptide database (humanPSMDb.txt.gz).** The tab-delimited file contains the following columns: peptide sequence, peptide mass, an indication of whether the peptide is a target or decoy, and a comma-separated list of the IDs of proteins containing this peptide. Dynamic modifications are indicated using bracket notation. Static modifications are not indicated. Decoy peptides do not have protein IDs associated with them.
3. **Ocean metaproteome peptide database (oceanPSMdb.txt.gz).** The tab-delimited file contains the following columns: peptide sequence, peptide mass, an indication of whether the peptide is a target or decoy, and a comma-separated list of the IDs of proteins containing this peptide. Dynamic modifications are indicated using bracket notation. Static modifications are not indicated. Note that none of the peptides have protein IDs associated with them, because the peptides came from shotgun sequencing data. Therefore, instead of the protein IDs, we subsitute the entire sequence from which the peptide was derived.
4. ***Plasmodium* peptide database (plasmodiumPSMDb.txt.gz).**The tab-delimited file contains the following columns: peptide sequence, peptide mass, an indication of whether the peptide is a target or decoy, and a comma-separated list of the IDs of proteins containing this peptide. Dynamic modifications are indicated using bracket notation. Static modifications are not indicated. Decoy peptides do not have protein IDs associated with them.
5. **PSMs (psmFile.txt.gz).** The tab-delimited file contains the following columns: name of the file that the spectrum resides in, scan number, charge, precursor m/z, and then for each search method the score and peptide sequence.
6. **R script 1 (createFinalFile.R).** This file contains the R script for creating the PSMs file (Supplemental File 2). The figures from this publication were generated from this file.
7. **R script 2 (rankingCurveFromFinalFile.R).** This file contains the R script for generating the figures in this publication.
8. **Commands (commands.txt).** Text file containing the database search commands, for various search engines, used to search the *E. coli* sample against the *E. coli* peptide database.

